# Differences in perceived severity of Zika virus infection and dengue fever and its influence on mosquito control practices in malaysia, a dengue-endemic country

**DOI:** 10.1101/061622

**Authors:** Li Ping Wong, Haridah Alias, Nasrin Aghamohammadi, I-Ching Sam, Sazaly Abu Bakar

## Abstract

**Background:** It is important to study the concerns over the Zika virus (ZIKV) outbreak among the general public in dengue-endemic countries such as Malaysia, as both diseases are transmitted by the same vector species. Furthermore, investigation of public prevention measures for ZIKV is essential in order to identify the gaps in mosquito control practices. The aims of this study were to explore the differences in 1) the perception of severity towards ZIKV infection and dengue fever, and 2) mosquito control practices before and after the ZIKV outbreak was declared a Public Health Emergency of International Concern (PHEIC).

**Method:** Data were collected between February 2015 and May 2016 using a computer-assisted telephone interviewing system on a random sample of 567 people from the general Malaysian population aged above 18 years from randomly selected households.

**Results:** The median scale score for perceived severity of ZIKV was 3 (interquartile range [IQR] 1-5) versus 4 (IQR 3-5) for dengue (P<0.001). The majority perceived dengue as being more severe than ZIKV (41.6%). Having friends or acquaintances that had died from dengue was significantly associated with higher perceived severity of ZIKV than dengue (odds ratio [OR] 1.913 [95% confidence interval (CI) 1.032-3.547]). The scores for mosquito control practices before and after ZIKV was declared a PHEIC were similar, at 4 (IQR 3-5). Multivariate analysis revealed that participants with a higher score for perception of severity of ZIKV were more likely to report greater mosquito control practices after the declaration of the PHEIC (OR 1.822 [95% CI 1.107-2.998]).

**Conclusions:** The emerging ZIKV pandemic requires concerted efforts to enhance mosquito control practices among the Malaysian public. Efforts to improve public mosquito control practices should focus on enhancing the perception of the severity of ZIKV.

**Author summary:** Investigation of the public perception of the severity of the re-emergence of Zika virus (ZIKV) in Malaysia, a dengue-endemic country, is of immense importance. It is also vital to know whether the public has heightened their mosquito prevention practices after the declaration of ZIKV as a Public Health Emergency of International Concern (PHEIC). The aim of this study was to explore the differences in 1) the perception of severity towards ZIKV infection and dengue fever, and 2) mosquito control practices before and after the ZIKV outbreak was declared a PHEIC. Findings showed that the public has a lower perception of severity of ZIKV than of dengue. Mosquito prevention practices were the same before and after the declaration of a PHEIC. People with a higher perception of severity of ZIKV reported higher mosquito control practices after the declaration of a PHEIC. The emerging ZIKV pandemic requires concerted efforts to enhance mosquito control practices among the Malaysian public. Efforts to improve public mosquito control practices should focus on enhancing the perception of severity of the ZIKV.

## Introduction

The current explosive pandemic re-emergence of mosquito-borne Zika virus infection (ZIKV) is causing worldwide concern. As ZIKV is typically transmitted by the *Aedes aegypti* mosquito, which also transmits dengue virus in Malaysia, there is tremendous anxiety among health authorities as the country already has a high number of dengue cases. Dengue incidence in Malaysia has continued to increase from 32 to 361 cases per 100,000 people between 2000 and 2014 [1]. Apart from sharing the same vector, ZIKV and dengue both result in high fever, but dengue can lead to haemorrhagic fever, which is potentially deadly. While dengue complications are far more serious for adults, ZIKV in contrast is much more dangerous for foetuses. ZIKV is commanding extensive media attention because of an alarming connection between ZIKV-infected women and microcephaly, a neurological disorder that results in babies being born with abnormally small heads and sometimes death [2,3]. The virus has been linked to over 4,000 cases of microcephaly in Brazil [4].

The Malaysian Ministry of Health put the country on alert following the declaration of ZIKV as a Public Health Emergency of International Concern (PHEIC) on 1^st^ February 2016 by the World Health Organization (WHO) [5]. All necessary precautions have been taken to limit the introduction of ZIKV from affected countries. Guidelines were issued for Malaysians to protect themselves against the ZIKV. Pregnant women, in particular, were advised to postpone travel to 24 countries in Central and South America where ZIKV has been detected. There is active surveillance in health clinics and hospitals for infected patients or cases of microcephaly. All levels of society have been urged to play their part in eliminating mosquito breeding sites.

To date, ZIKV infection has not been reported in Malaysia since an early description in 1966 [6]. Little is known of perceptions of the general Malaysian public towards the ZIKV outbreak. How seriously a person perceives a disease is important as it has a direct association with many important health outcomes such as their level of functioning and ability, utilization of health care and adherence to treatment plans laid out by health-care professionals [7]. A previous nationwide study of dengue prevention practices in Malaysia revealed that public perceived severity of dengue fever was a significant factor associated with higher prevention practices [8]. It is hypothesized that the perceived severity of ZIKV likewise may also be associated with enhanced mosquito control practices. Little is known about the effect of the PHEIC declaration on mosquito prevention and control practices of the Malaysian public

The aim of this study, therefore, is to assess differences in 1) the perception of severity towards ZIKV infection and dengue fever, and 2) mosquito control practices before and after the declaration of the ZIKV outbreak as a PHEIC. The study firstly hopes to uncover factors that prompt the public to heighten their mosquito control practices in light of the new emergence of ZIKV and secondly whether or not the perceived severity of ZIKV influences mosquito control practices. The findings will have importance for public policymakers in enacting new policies that promote mosquito control practices now that *Aedes aegypti* mosquitoes are spreading both dengue and ZIKV.

## Methods

### Sample

Interviews were conducted between February 2015 and May 2016 using a computer-assisted telephone interview (CATI) system. Sampling was accomplished by random digit-dialling of landline phone numbers from all the 11 states and two federal territories in Peninsular Malaysia. Only one participant per household was randomly selected to take part in the survey. Eligible participants were 18 years of age or older and had heard of ZIKV. Interviews were conducted between 5.30 p.m. and 10.00 p.m. on weekdays and from 12.00 p.m. to 7.00 p.m. at weekends or during public holidays to avoid over-representation of unemployed participants. Unanswered calls were attempted at least twice more on separate days before being regarded as non-responses.

### Instrument

The questionnaire was divided into five sections. The first and second sections ascertained the participants’ socio-demographic background, their surrounding environment and dengue experiences. The third section investigated the differences between the perceived severity of ZIKV and dengue fever. Participants were asked: “On a scale of 0 (not worried at all) to 6 (worried all the time), how worried are you about ZIKV versus dengue fever?” The response options comprised the categories “not at all/rarely/occasionally/sometimes/frequently/usually/all the time”, and were scored 0, 1, 2, 3, 4, 5 and 6, respectively, with higher scores representing a higher perception of severity. A higher score for perception of severity of ZIKV compared to dengue fever was used as the dependent variable in the multivariate logistic regression model.

The fourth section determined differences in mosquito control practices before and after the ZIKV outbreak was declared a PHEIC. The response options were the same categories as above, with higher scores representing a higher level of mosquito control practices. The dependent variable in the multivariate logistic regression model was a higher score for mosquito control practices after compared to before the declaration of a PHEIC. The questionnaire is provided in Appendix 1.

The questionnaires were in three languages: Bahasa Malaysia (the national language of Malaysia), English and Chinese. The content of the questionnaire was validated by a panel of experts and pilot-tested prior to data collection. A team of trained interviewers from different ethnic groups conducted the interviews; each interviewer was assigned to interview respondents of a similar ethnic group in their native languages. Informed consent was obtained verbally. Respondents were assured that their responses would be confidential and reminded that their participation was voluntary. The study was approved by the Medical Ethics Committee, University Malaya Medical Centre, Kuala Lumpur, Malaysia (MECID NO: 20162-2194).

### Data analyses

Descriptive analysis was performed to determine the frequency distribution of demographic factors, perception of severity and mosquito control practices. Perception of severity and mosquito control practices were defined as dependent variables and measured on a Likert scale, and presented as median and interquartile range (IQR). Univariate analysis was first used to determine the associations between independent and dependent variables. Multivariate logistic regression analyses were used to investigate factors associated with 1) a higher level of perceived severity of ZIKV infection than dengue, and 2) higher mosquito control practices after than before the declaration of the ZIKV outbreak as a PHEIC. All variables with p<0.05 in the univariate analysis were included in multivariate logistic regression analyses using the “enter” method. Odds ratios (OR), 95% confidence intervals (95% CI) and P-values were calculated for each independent variable. The model fit was assessed using the Hosmer-Lemeshow goodness-of-fit [9]. All analysis was performed with IBM SPSS Statistics version 20.0 (IBM, USA).

## Results

### Socio-demographics, dengue experiences and surrounding environment

Figure 1 presents a flow chart of the CATI process. A total of 4,675 call attempts were made, resulting in 567 responding households. The response rate of 70.8% was computed as the number of completed interviews divided by the number of contacted and eligible households (n = 801). As shown in Table 1, the majority of the respondents were females (n = 403, 71.1%) and were ever married (n = 436, 76.9%). Most of the respondents were Malay (n = 426, 75.1%). The majority had a secondary education (n = 306, 54.0%). By occupation, most of the respondents were housewives (n = 169, 29.8%), or of the professional and managerial category (n = 167, 29.5%). The majority of respondents reported having an average monthly household income of RM2001 to RM4000 (one Malaysian ringgit is equal to USD 0.25). Out of the total study sample, only 9.3% (n = 53) had experience of having dengue fever. Approximately half of the participants noted that dengue was a problem in their neighbourhood (n = 292, 51.5%).

**Fig. 1.**
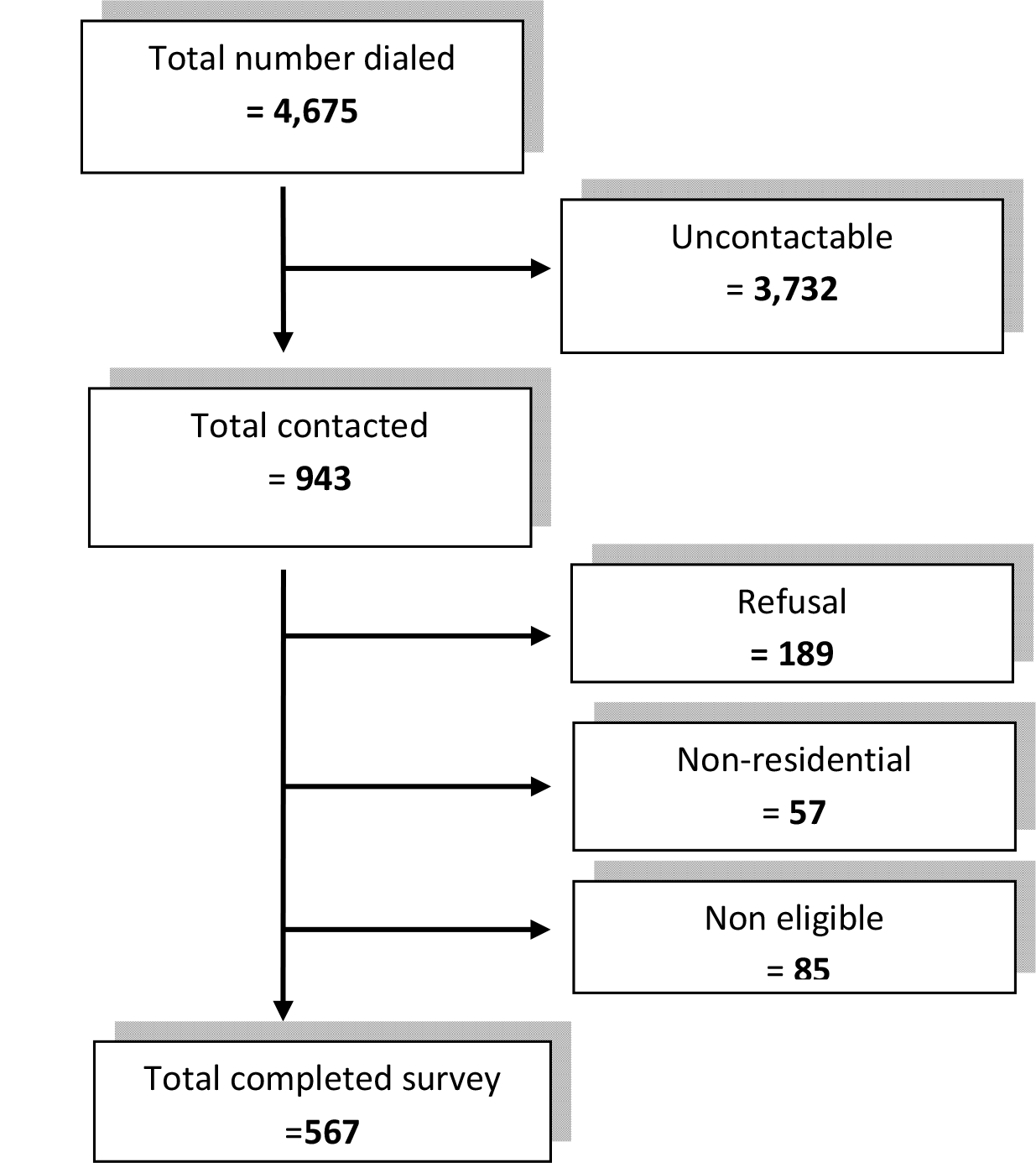
CATI process flowchart.

**Table 1.**
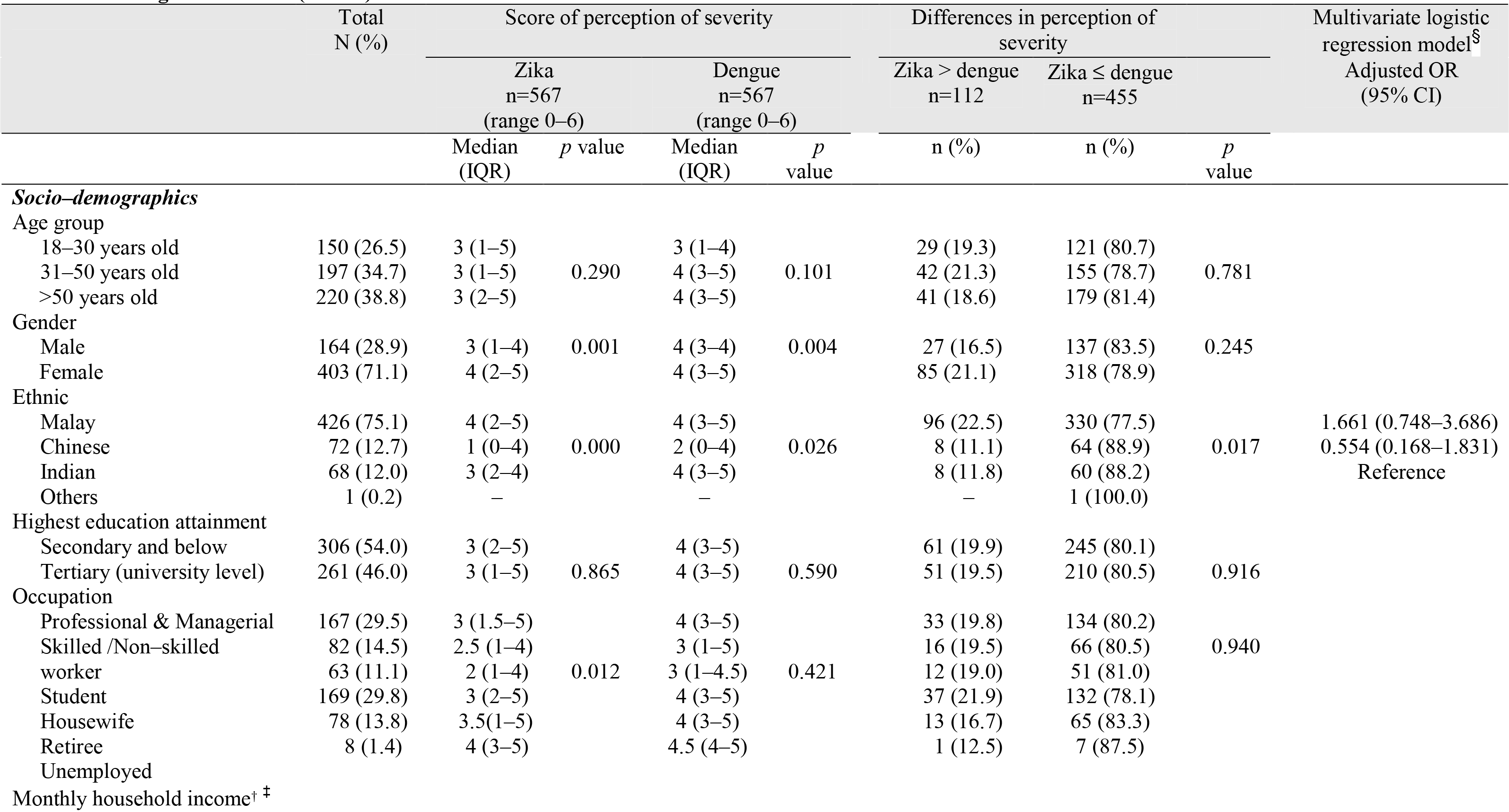
Perception of severity between Zika virus infection and dengue fever in general by socio-demographic characteristics, dengue experience and surrounding environment (N=567)

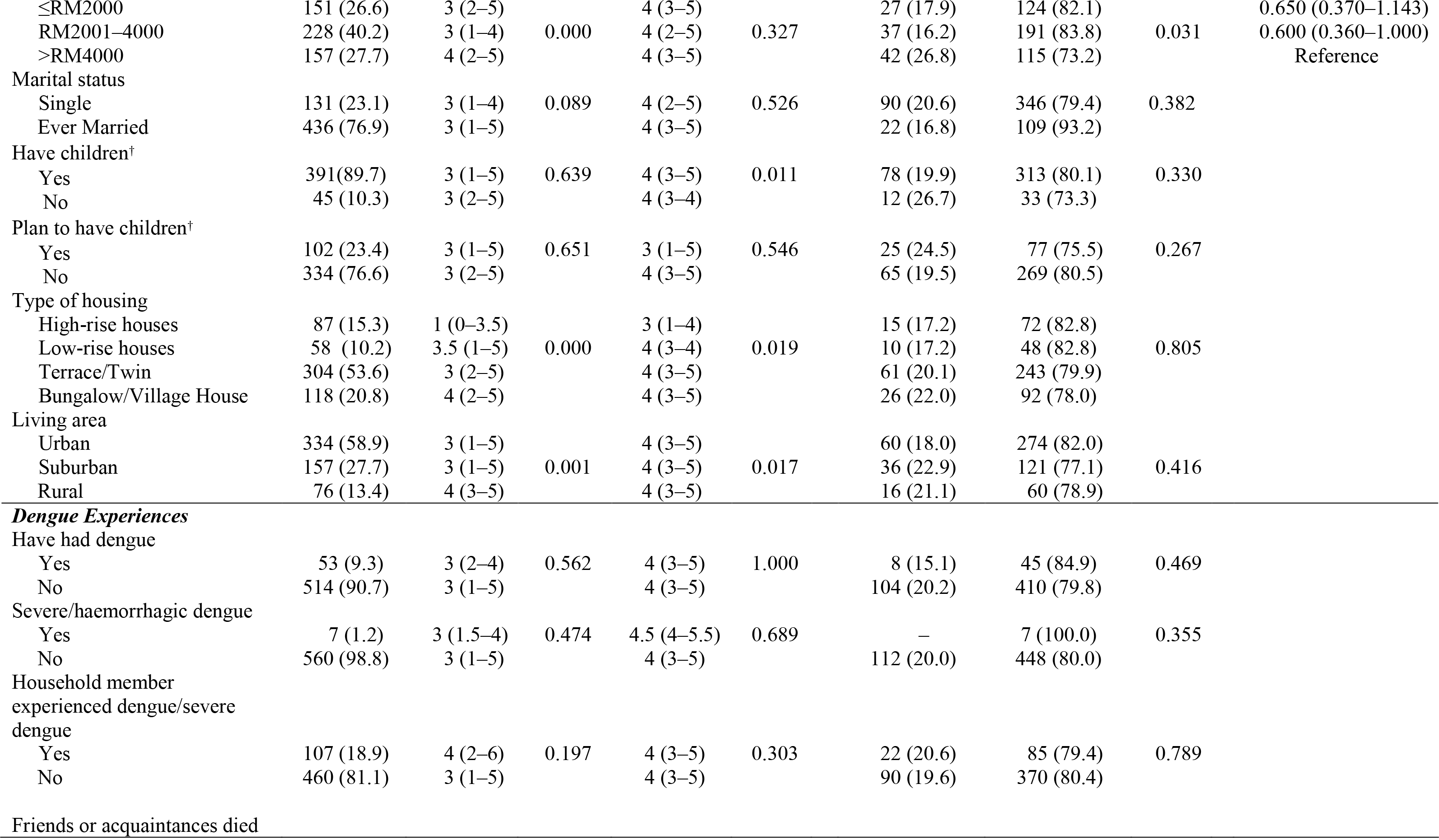

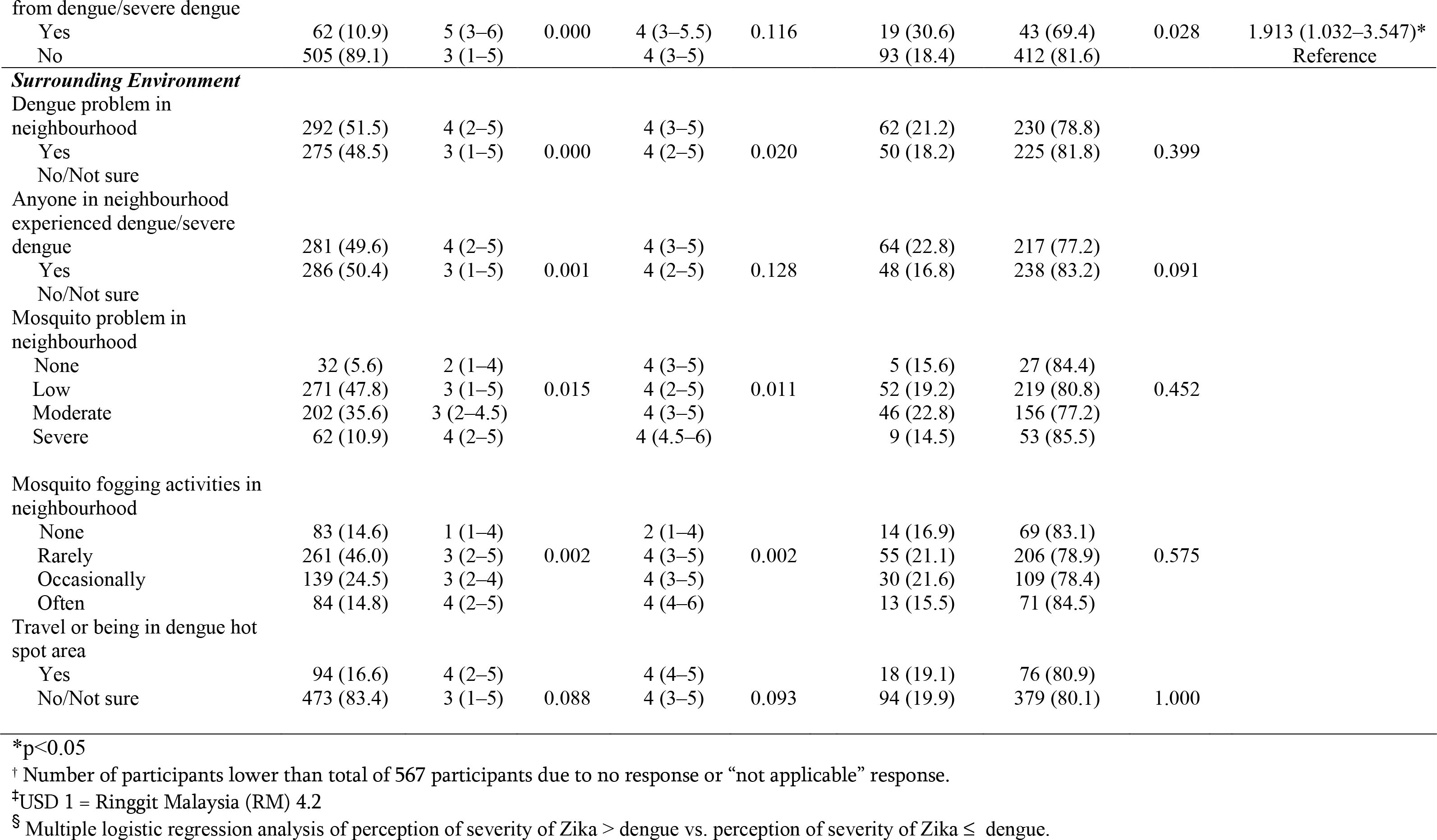

### Perception of severity towards ZIKV and dengue fever

Table 1 shows that the overall participants’ median score for the perception of severity of ZIKV was 3 (IQR 1-5), which was significantly lower than that of dengue fever (4 [IQR 3-5], p<0.001). The median scores for the perception of severity of ZIKV was significantly higher the following groups, with respect to their comparator groups: females, household income >RM4000, living in a bungalow or single village house, residence in a rural area, having friends or acquaintances who had died from dengue, reporting dengue as a problem in their neighbourhoods, having a neighbour with dengue fever, increasing perception of a mosquito problem in their neighbourhood, and increased frequency of fogging in the neighbourhood.

### Differences in perceptions of severity towards ZIKV infection and dengue fever

The majority of the participants perceived dengue infection as being more severe than ZIKV infection (41.6%, n = 236). The proportion of participants with a similar perception of severity for both ZIKV and dengue was 38.6% (n = 219). Only 19.8% (n = 112) perceived ZIKV infection to be more severe than than dengue fever, and by univariate analysis, these were more likely to be participants of Malay ethnicity, those with a monthly household income >RM4000, and those with friends or acquaintances who had died from dengue. In multivariate analysis, only having friends or acquaintances who had died from dengue was significantly predictive of a higher perception of severity for ZIKV infection than dengue fever (OR 1.913 [95% CI 1.032-3.547]). The final model accounted for 26.0% of the total variability in perception of severity of ZIKV (R^2^ = 0.260), and the Hosmer-Lemeshow test was nonsignificant (Χ^2^ = 1.730, *P* = .943), indicating good model fit.

### Mosquito control practices after the declaration of the PHEIC

Table 2 summarizes the analyses of scores of mosquito control practices. The overall participants’ median (IQR) scores for mosquito control practices before and after ZIKV was declared a PHEIC were similar, at 4 (IQR 3-5). With respect to their comparator groups, participants from the following groups recorded the highest median scores: >50 years, non-working participants (housewives, students and unemployed) compared to working participants, low monthly household income groups, married participants, having a household member who had experienced dengue, having friends or acquaintances who had died from dengue, and a higher perception of severity for ZIKV than dengue. Notably, there was no significant difference in mosquito control practice scores after the declaration of a PHEIC between participants with children and without children, and surpristingly, participants who plan to have children reported a significantly lower median mosquito control practice score than those who do not plan to have children. With regard to type of housing, the median mosquito control practice score was significantly lower among participants living in high-rise houses.

**Table 2.**
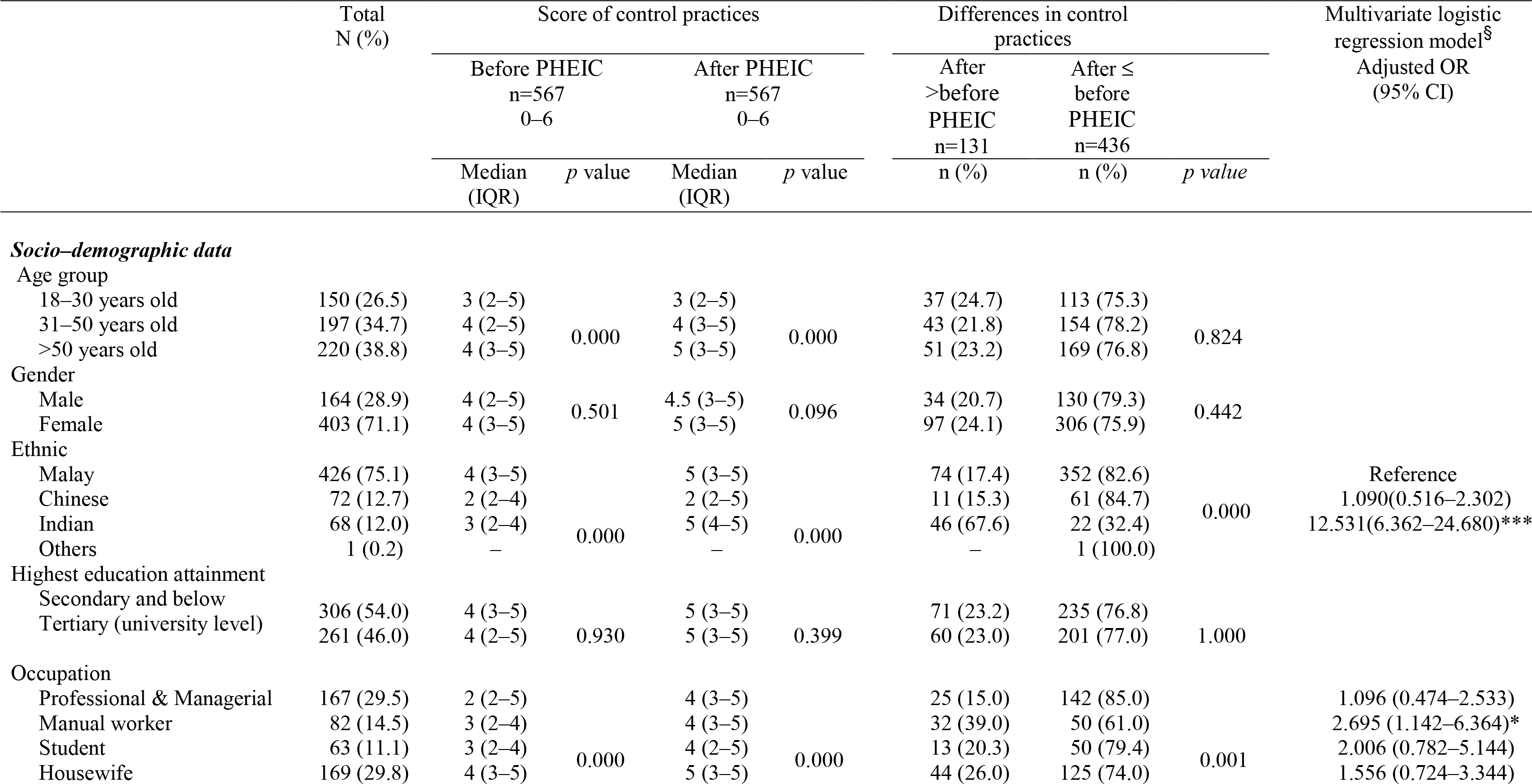
Mosquito control practices before and after ZIKV was declared a PHEIC by socio-demographic characteristics - dengue experience and surrounding environment

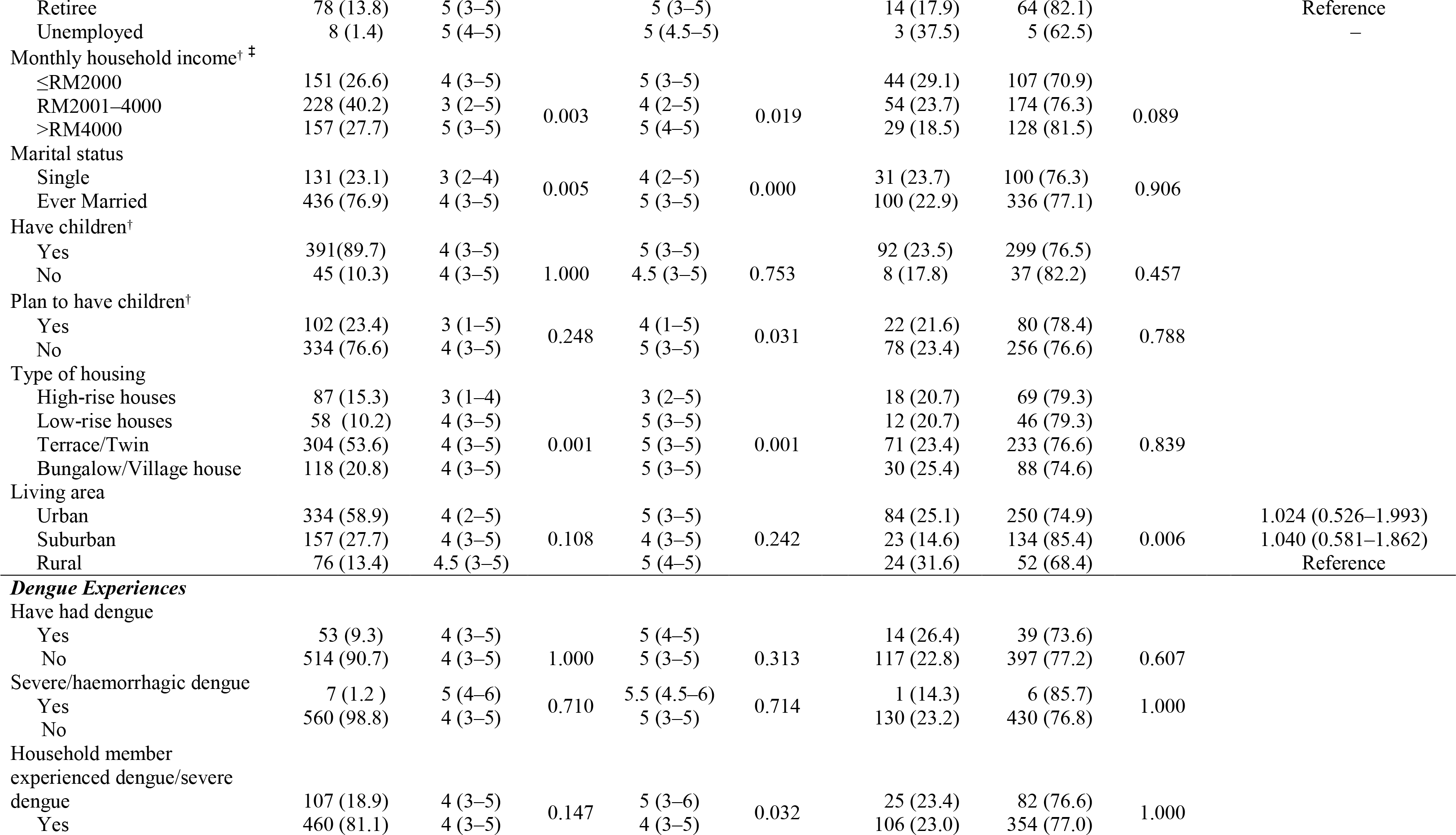

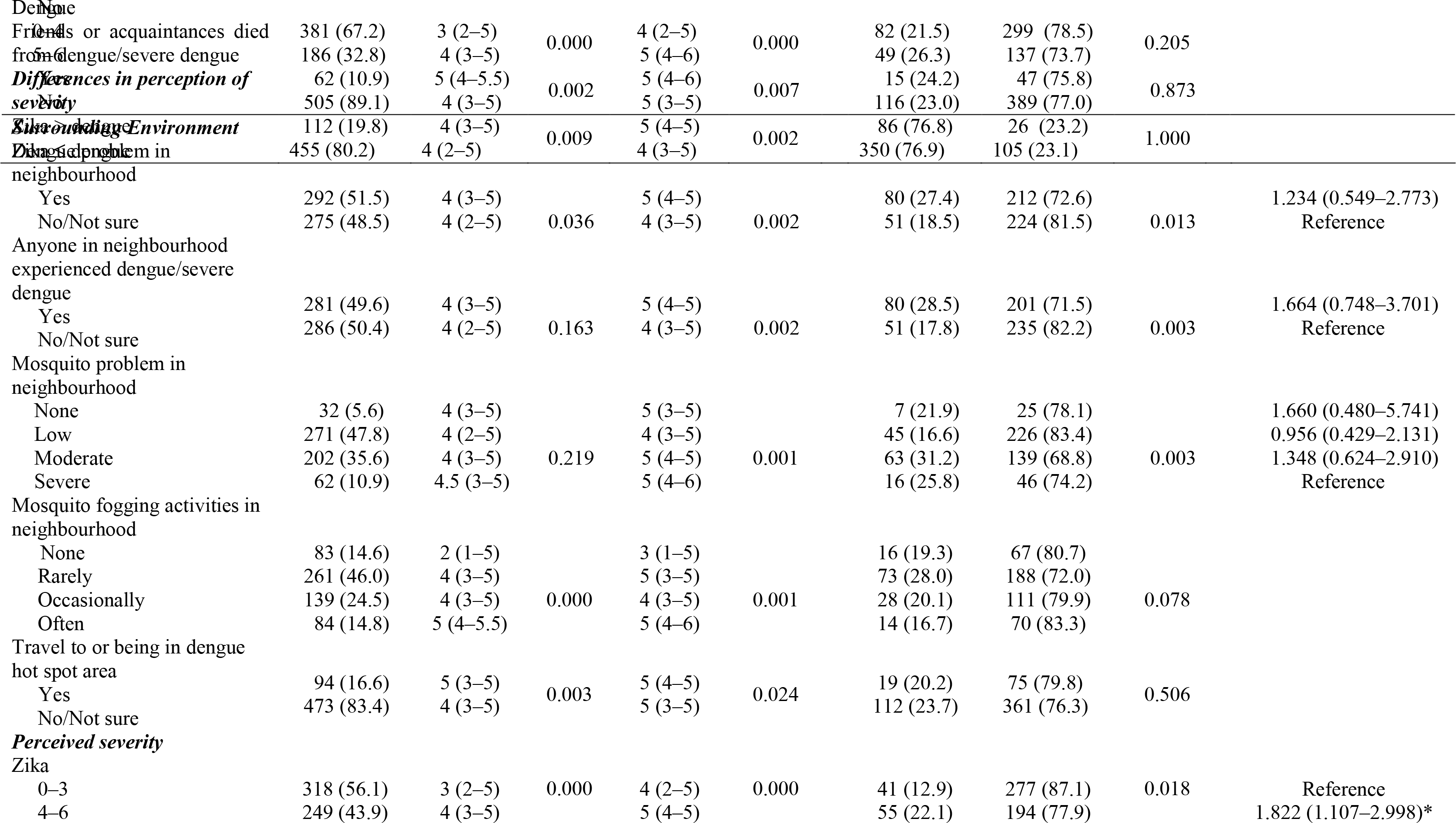

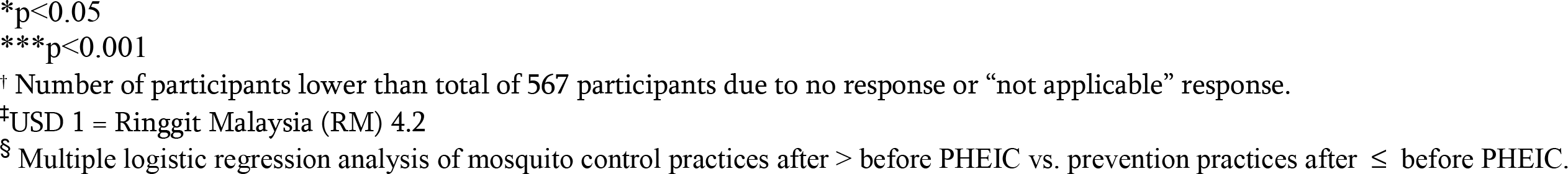

In terms of the surrounding environment, participants with a higher median score for mosquito control practices were more likely to have reported dengue as a problem in their neighbourhoods, have a neighbour with dengue fever, be from areas with severe and moderate mosquito problems, have frequent mosquito fogging activities in their neighbourhoods, and have traveled to or live in a dengue hot spot area. Higher scores in the perceived severity of both ZIKV and dengue fever were significantly associated with higher median mosquito control practice scores both before and after the declaration of a PHEIC.

### Differences in mosquito control practices before and after the declaration of the PHEIC

Approximately 23% (n = 131) of participants reported higher mosquito control practices after compared to before the declaration of a PHEIC, and in univariate analysis, this was associated with ethnicity, occupational groups, area of residence, living in a neighbourhood with a dengue problem, having a neighbour who had experienced dengue, living in a neighbourhood with a moderate or severe mosquito problem, and those with a higher perceived severity of ZIKV score of 4-6. All seven of these variables were included in the multivariate analysis. Having a score of perceived severity of ZIKV of 4-6 (OR 1.822 [95% CI 1.107-2.998]), Indian ethnicity (OR 12.531 [95% CI 6.362-24.680]) and manual workers (OR 2.695 [95% CI 1.142-6.364]) remained as significant independent predictors of higher mosquito control practices after compared to before the ZIKV outbreak was declared a PHEIC. The final model accounted for 23.4% of the total variability in mosquito control practices (R^2^ = 0. 234), and the Hosmer-Lemeshow test was nonsignificant (χ^2^ = 6.546, *P* = .586), indicating good model fit.

## Discussion

Despite the massive ongoing media coverage in Malaysia about the ZIKV epidemic, our study found that the level of perceived severity of ZIKV among the study participants is moderate, with a median score of 3 out of a possible 6. Dengue fever, in contrast, has been a longstanding problem in Malaysia, and the median score for the perceived severity of dengue was 4, which was just over the midpoint. This implies that the overall perception of the severity of both ZIKV and dengue is not high among the study population, with the perception of severity for dengue being slightly higher than for ZIKV.

Only approximately 20% of the study participants rated the perceived severity of ZIKV higher than that of dengue. There may be several reasons for this. Firstly, as ZIKV was unheard of among the general Malaysian public, and to date, no cases have been reported in Malaysia since the recent ZIKV outbreaks, the majority may not view the outbreak as severe as ZIKV has not affected Malaysia. If so, this implies the need to build community preparedness for potential ZIKV outbreaks here, which will heighten the perception of risk of this re-emerging infectious disease and may bring community attention to the need to strengthen the public health response infrastructure [10]. Secondly, many may be aware that fatality from ZIKV infection is rare, in contrast to dengue, where the possibility of deadly complications from severe dengue is higher.

Therefore, for individuals who have no plans to become pregnant or are not currently pregnant, ZIKV infections may not be of much concern to them. In addition, it has been publicized in the media that ZIKV symptoms are usually mild compared to dengue. The main consequences of ZIKV infection are severe foetal brain defects [11,12], therefore this could be why the majority perceived the severity of dengue to be higher than that of ZIKV. Nevertheless, further investigation into the reasons for such a low perception of severity of ZIKV would be worthwhile to provide evidence-based insights into future interventions to enhance the perception of severity.

It would be expected that ZIKV would most concern pregnant women, or couples who are planning to have children. However, it is a worrying finding in the study that neither married participants nor those who plan to have children have a high perception of severity of ZIKV. Further investigation is warranted to find out whether this is due to a lack of knowledge or awareness or other reasons. Women who are planning to become pregnant and their partners should be targets of intervention to increase awareness of the serious health effects of ZIKV on unborn babies.

Although univariate results revealed that those who reported mosquitoes and dengue as a problem in their neighbourhood were more likely to have higher perception of severity of ZIKV, communities less affected by dengue should also be a target of intervention to increase perception of severity of ZIKV. The multivariate analysis of factors associated with a higher perceived severity of ZIKV found that knowing friends or acquaintances who had died from dengue was the only significant predictor. The resulting experience or trauma may be associated with a stronger feeling of concern over a new mosquito-borne disease.

A worrying finding is that the scores of mosquito control practices before and after the declaration of a PHEIC were similar, and only 23% reported higher mosquito control practices after the declaration. Although the ZIKV outbreak generated considerable local and global media attention, it did not appreciably change the general public’s mosquito control efforts. It was notable that participants from neighbourhoods with dengue and mosquito problems and those with a neighbour with dengue fever reported higher mosquito prevention practices since the declaration of the PHEIC. Multivariate analysis identified a higher score of perceived severity of ZIKV infection as a determinant of higher mosquito control practices, which indicates the importance of influencing public perception of severity of ZIKV to increase mosquito prevention practices.

## Strengths and limitations

The main strength of this study is the use of a population-based landline sample with a good response rate over a broad age range. The demographics of the participating population are similar to the general population, suggesting that these results may be reflective of national trends. However, it is acknowledged that this study has several limitations. Firstly, the limitation of telephone surveys includes a lack of representativeness of households with no landline telephone. However, the use of mobile phones does not allow stratification by geographical region. Another limitation is the possibility of self-reporting response bias.

## Conclusions

This study aimed to gauge public perception of the severity of the consequences of ZIKV infection and the impact on the public’s mosquito control practices. Findings revealed a moderate perception of severity of ZIKV. The perception of severity of dengue was higher than that of the ZIKV. Having friends or acquaintances who had died from dengue was the most important contributing factor to the higher perception of ZIKV compared to dengue. There was no increase in mosquito control practices before or after the ZIKV outbreak was declared a PHEIC. A higher perceived severity of ZIKV was the only significant independent predictor of having higher mosquito prevention practices. These results provide evidence of the need to enhance mosquito control practices among the Malaysian public by enhancing the public perception of the severity of ZIKV.

## References

1. Mudin RN. Dengue incidence and the prevention and control program in Malaysia. International Medical Journal of Malaysia. 2015 Jun 1;14(1):05–10.

2. Cauchemez S, Besnard M, Bompard P, Dub T, Guillemette-Artur P, Eyrolle-Guignot D, Salje H, Van Kerkhove MD, Abadie V, Garel C, Fontanet A.Association between Zika virus and microcephaly in French Polynesia, 2013-15: a retrospective study. Lancet. 2016 May 27;387(10033):2125–32. doi: 10.1016/S0140-6736(16)00651-6

3. Schuler-Faccini L. Possible association between Zika virus infection and microcephaly: Brazil, 2015. MMWR. Morbidity and Mortality Weekly Report. 2016 Jan 29;65(3):59–62

4. Fauci AS, Morens DM. Zika virus in the Americas: yet another arbovirus threat. New England Journal of Medicine. 2016 Jan 13;374:601–604. doi: 10.1056/NEJMp1600297

5. Sam JIC, Chan YF, Vythilingam I, Wan Sulaiman WY. Zika virus and its potential re-emergence in Malaysia. Med J Malaysia 2016; 71 (2): 68–70.

6. Marchette NJ, Garcia R, Rudnick A. Isolation of Zika virus from *Aedes aegypti* mosquitoes in Malaysia. Am J Trop Med Hyg 1969; 18(3): 411–5.

7. Petrie KJ, Weinman J. Patients’ perceptions of their illness. The dynamo of volition in health care. Current Directions in Psychological Science. 2012 Feb 1;21(1):60–5. doi: 10.1177/0963721411429456

8. Wong LP, Shakir SM, Atefi N, AbuBakar S. Factors affecting dengue prevention practices: nationwide survey of the Malaysian public. PLoS One. 2015 Apr 2;10(4):e0122890. doi:10.1371/journal.pone.0122890

9. Hosmer Jr DW, Lemeshow S, Sturdivant RX. Model-building strategies methods for logistic regression. Applied Logistic Regression, 3^rd^ edition. 2000:89–151

10. Katz A, Staiti AB, McKenzie KL. Preparing for the unknown, responding to the known: communities and public health preparedness. Health Affairs. 2006 Jul 1;25(4):946–957. doi: 10.1377/hlthaff.25.4.946

11. Driggers RW, Ho CY, Korhonen EM, Kuivanen S, Jaaskelainen AJ, Smura T, Rosenberg A, Hill DA, DeBiasi RL, Vezina G, Timofeev J. Zika virus infection with prolonged maternal viremia and fetal brain abnormalities. New England Journal of Medicine. 2016 Mar 30;374:2142–2151. doi: 10.1056/NEJMoa1601824

12. Nunes ML, Carlini CR, Marinowic D, Neto FK, Fiori HH, Scotta MC, Zanella PL, Soder RB, da Costa JC. Microcephaly and Zika virus: a clinical and epidemiological analysis of the current outbreak in Brazil. Jornal de Pediatria (Versao em Portugues). 2016 Jun 3;92(3):230–40. doi: 10.1016/j.jped.2016.02.009. Epub 2016 Apr 15.

